## Supplementary file 1 for "Layers of immunity: Deconstructing the *Drosophila* effector response"

**Supplementary fly genetics considerations**

The process to generate these combinatorial mutations required recombination, introgression, and the combining of chromosomes. Below we highlight the key considerations that were taken in constructing these lines, and provide some advice on their use.

- Lines involving the *Hayan-psh^Def^* mutation on the X chromosome are ΔTOLL, ΔMel in genotype. This mutation is surprisingly well-tolerated, can be kept homozygous, and requires no special care. A rapid phenotypic screen to validate the presence of the mutation can be done by tearing a larva open with forceps for a melanization assay. This can be done on filter paper, or in a small well (e.g. microtube, cavity microscope slide). While wild-type individuals will blacken over 2-3 minutes due to serine protease activity catalyzing the melanization reaction, *Hayan-psh^Def^* mutant larvae fail to melanize for >30 minutes. The melanization reaction eventually takes place regardless, as it occurs naturally in the absence of serine proteases and the phenoloxidases which they activate.
- Lines involving the *spz^rm7^* mutation are ΔTOLL and homozygous sterile. There can sometimes be difficulty with obtaining good homozygous numbers with these mutants. In personal communications with others using this line, this difficulty to obtain homozygous individuals may be lab-specific (e.g. local microbiota, food recipe, etc…). Rearing individuals at low density can encourage higher homozygous numbers. This is particularly important to consider In combination with the *PPO1, PPO2* double mutation, which produces very few homozygotes – anecdotally, at times as little as a ~1/1000 homozygous rate.
- Lines involving the *Rel^E20^* mutation are ΔIMD, and are relatively straightforward to rear homozygous in our hands. This genotype has a slightly delayed development (~1 day) compared to wild-type flies. Due to the severe immunodeficiency of this line, and deficiency in certain physiological processes such as epithelial gut renewal, poor rearing conditions (e.g. long periods between flipping library stocks) may lead to crashes more easily than standard fly strains. However, we have had no difficulties maintaining this stock homozygous indefinitely on varying food recipes. In personal communications with others, sometimes keeping *Rel^E20^* over a third chromosome balancer is useful to prevent stochastic stock collapse.
- Lines involving Δ*PPO1,* Δ*PPO2* are ΔMel and again, relatively straightforward to keep. Like Rel^E20^, this strain can be more sensitive to periods of neglect. In addition, adult fecundity may depreciate slightly faster than other strains, and so we would recommend sticking closely to standard Drosophila rearing protocols that refresh adults regularly. Like Hayan-psh^Def^, a rapid phenotypic assay to verify the mutation is present is to tear a larva open and assess the speed of the melanization reaction. PPO1, PPO2 double mutant larvae fail to melanize for >30 minutes. The PPO1 mutation includes a white^+^ insertion, while the PPO2 mutation does not and must be screened by PCR; see Binggeli et al. (2014).
- Lines involving NimC1^1^; eater^1^ are ΔPhag, and both mutations bear a white^+^ insertion. NimC1^1^; eater^1^ flies were the most complex to work with in our hands. First, the w; NimC1^1^; eater^1^ strain originally used in Melcarne et al. (2019) presented no major issues in standard rearing. We have deposited this non-isogenic strain in the VDRC DrosDel Immunity Panel (<https://shop.vbc.ac.at/vdrc_store/drosdel-info>) for this reason.
  We encountered strange behaviours in various iterations of the isogenic NimC1, eater double mutant. Our original strain appeared to have a highly depressed IMD pathway response, which we could rescue somewhat, but not fully, with additional recombinations to the DrosDel background. At another point, we were forced to regenerate this line by combining the NimC1^1^ mutation (chromosome 2) and eater^1^ mutation (chromosome 3). Despite starting from the same source populations, this new line was homozygous lethal. We have no explanation for this fecundity behaviour. As a result, we were forced to complete certain experiments in the study with the non-isogenic w; NimC1^1^; eater^1^ strain, which behaved consistently with our previous data. We have since additionally recombined both the NimC1^1^ and eater^1^ mutations to the DrosDel background, and produced flies that produce homozygotes, but are homozygous sterile in the lab. This line can be maintained balanced as NimC1^1^/CyO and homozygous for eater^1^, providing a straightforward visual cue (curly wings) to select homozygous adults. In the future, combination with a GFP-labelled 2^nd^ chromosome balancer will be useful to enable study of larval hemocytes.

The parent balancer strains used to combine and recombine these mutations are as follows (“iso” = isogenic DrosDel chromosome):

- Fk/FM7; iso; iso
- iso; Gla/CyO; iso
- iso; iso; TM2/TM6c, sb

Standard *Drosophila* genetic crossing schemes were performed to combine the mutations above, with mutations typically collected over the CyO and TM6c balancer chromosomes.
