## Supplementary figures and images for "Layers of immunity: Deconstructing the *Drosophila* effector response"

### A. fumigatus NI.png

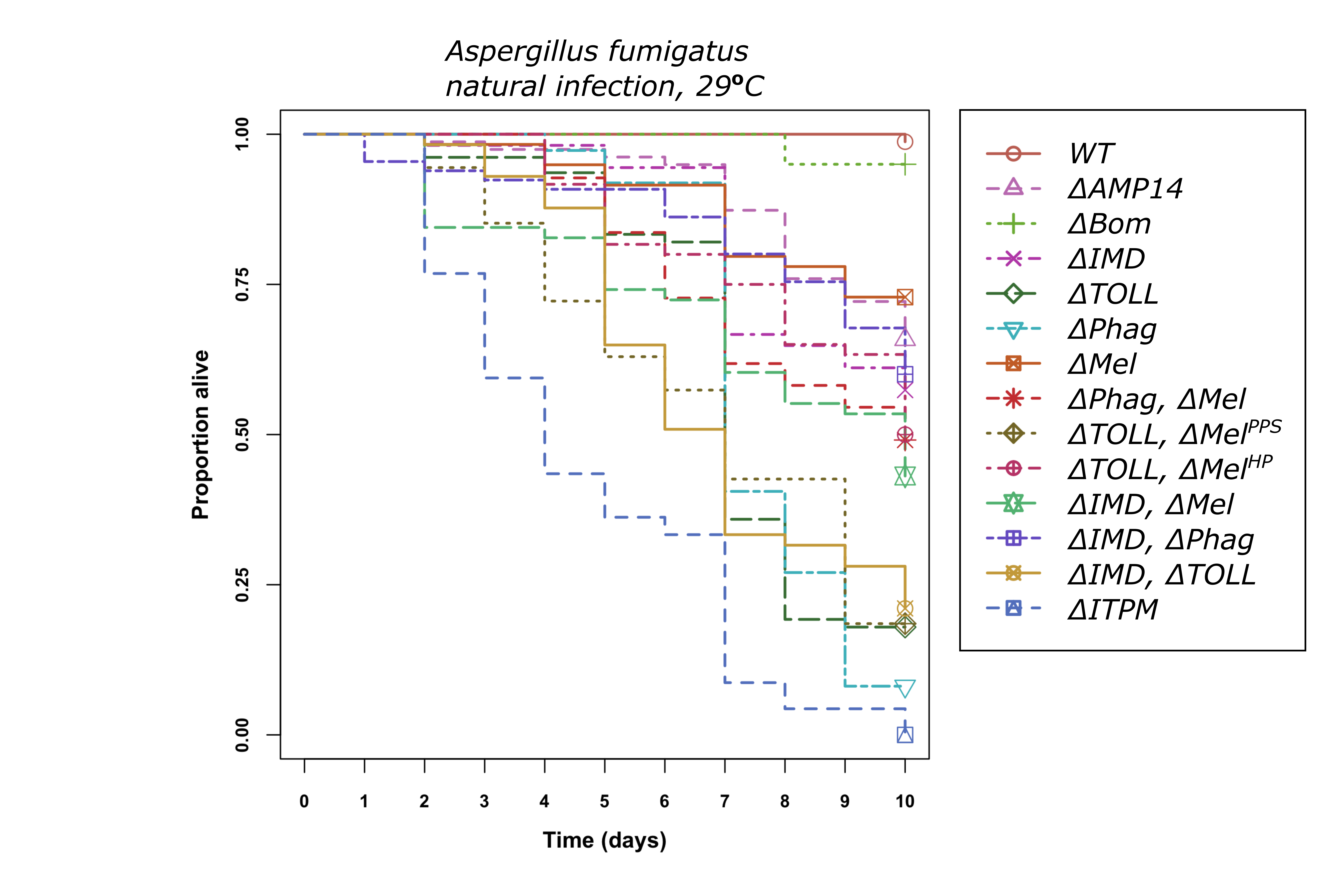

### B. bassiana NI.png

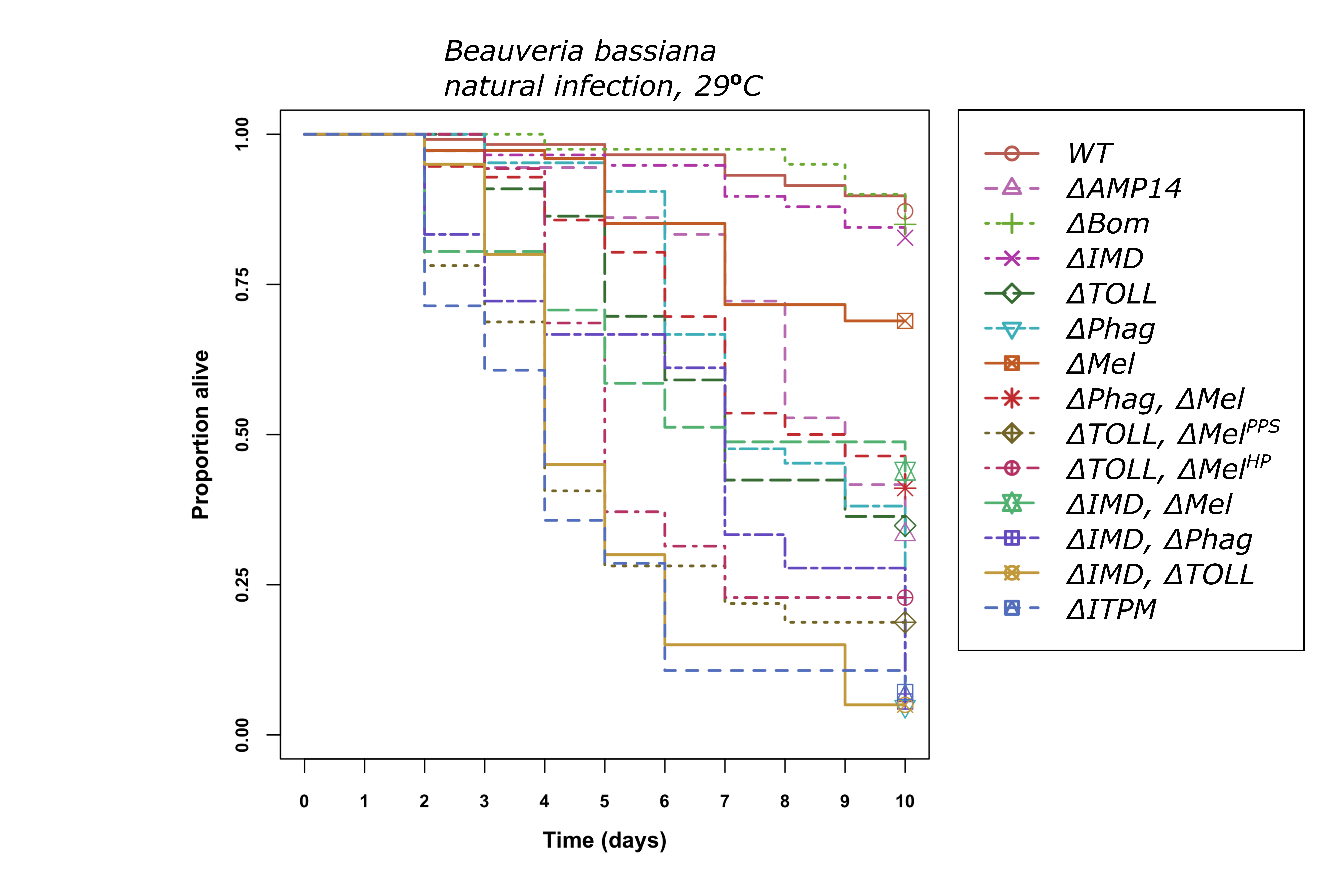

### B. bassiana SI.png

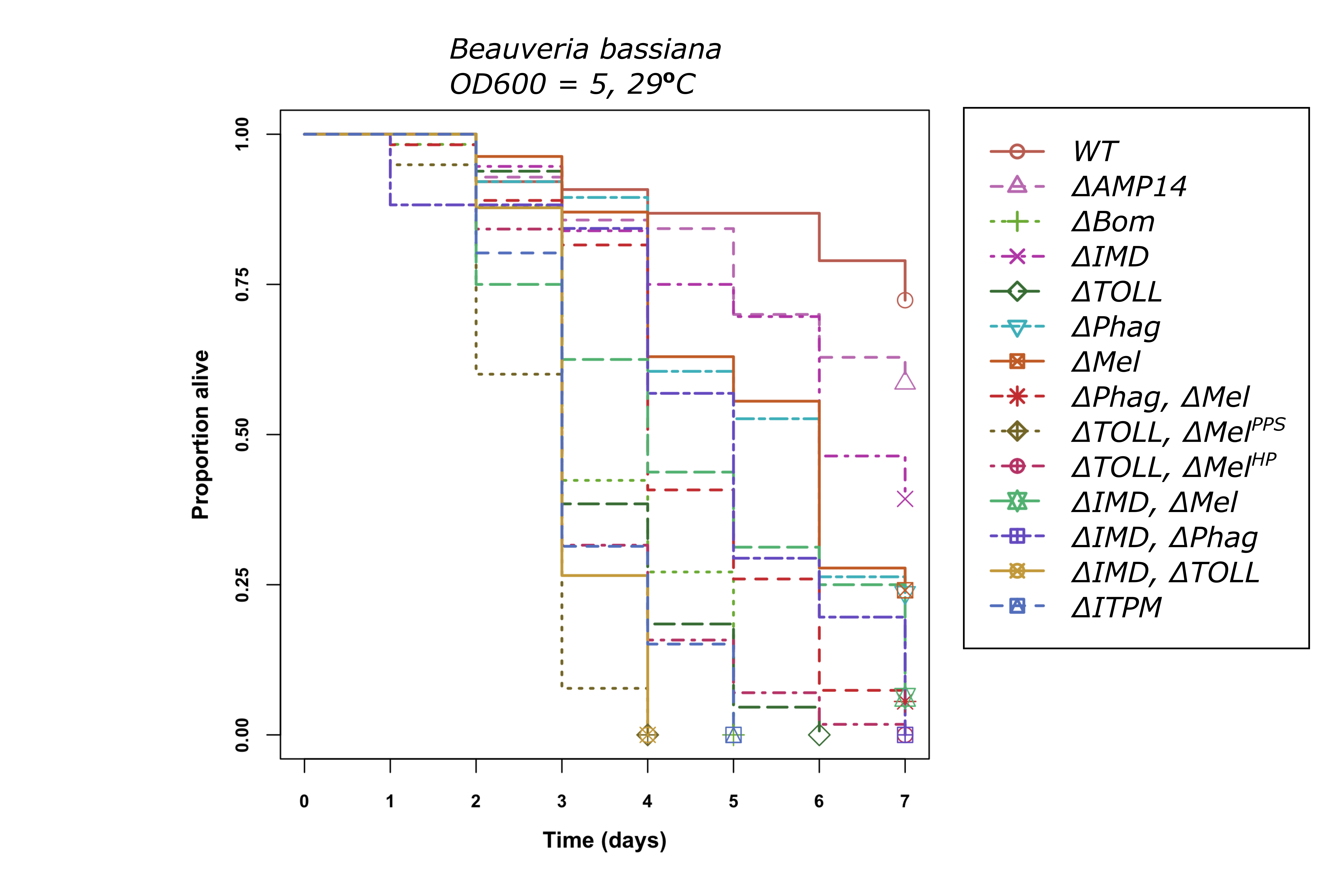

### B. subtilis.png

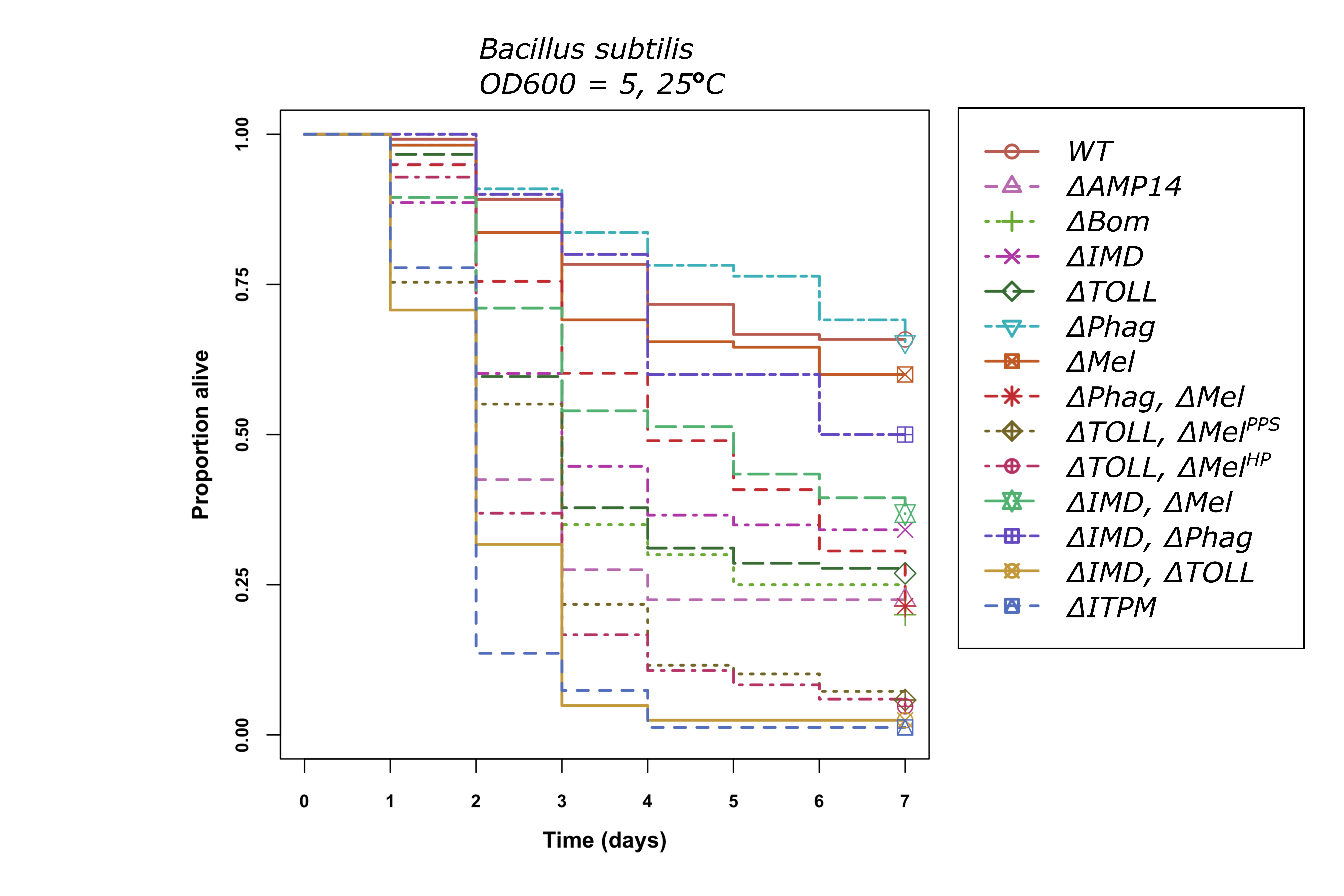

### C. albicans SI.png

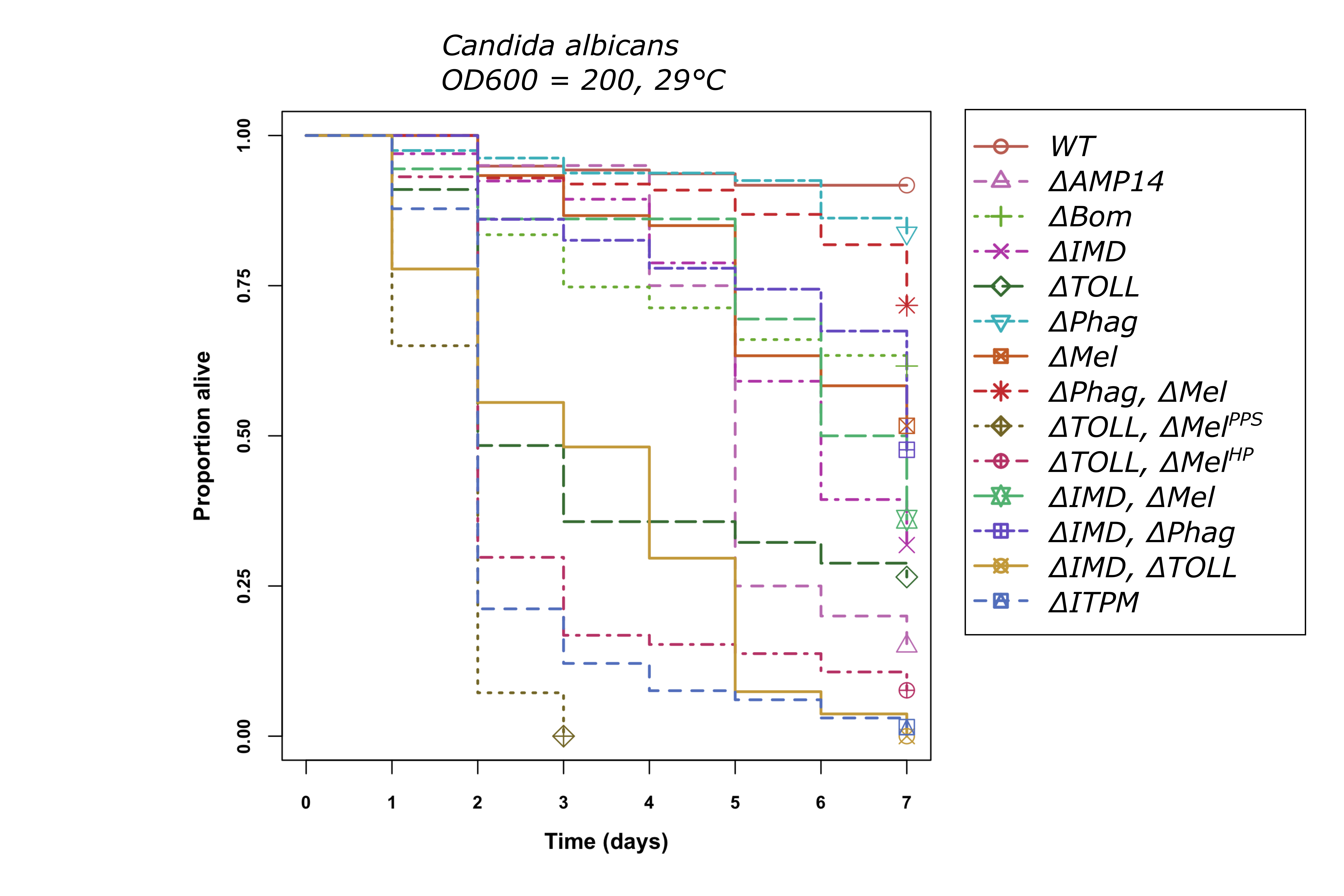

### C. diphtheriae.png

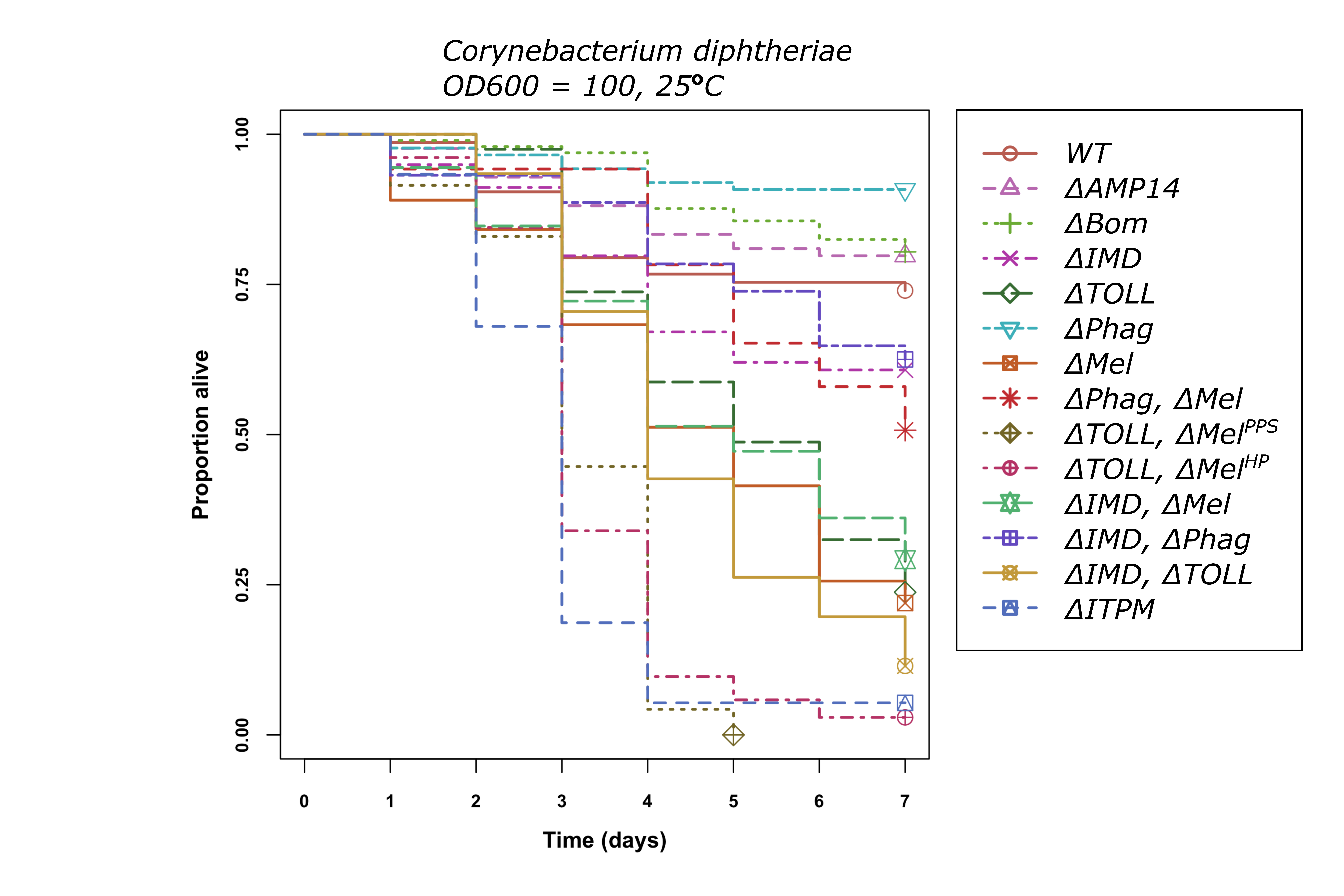

### DCV.png

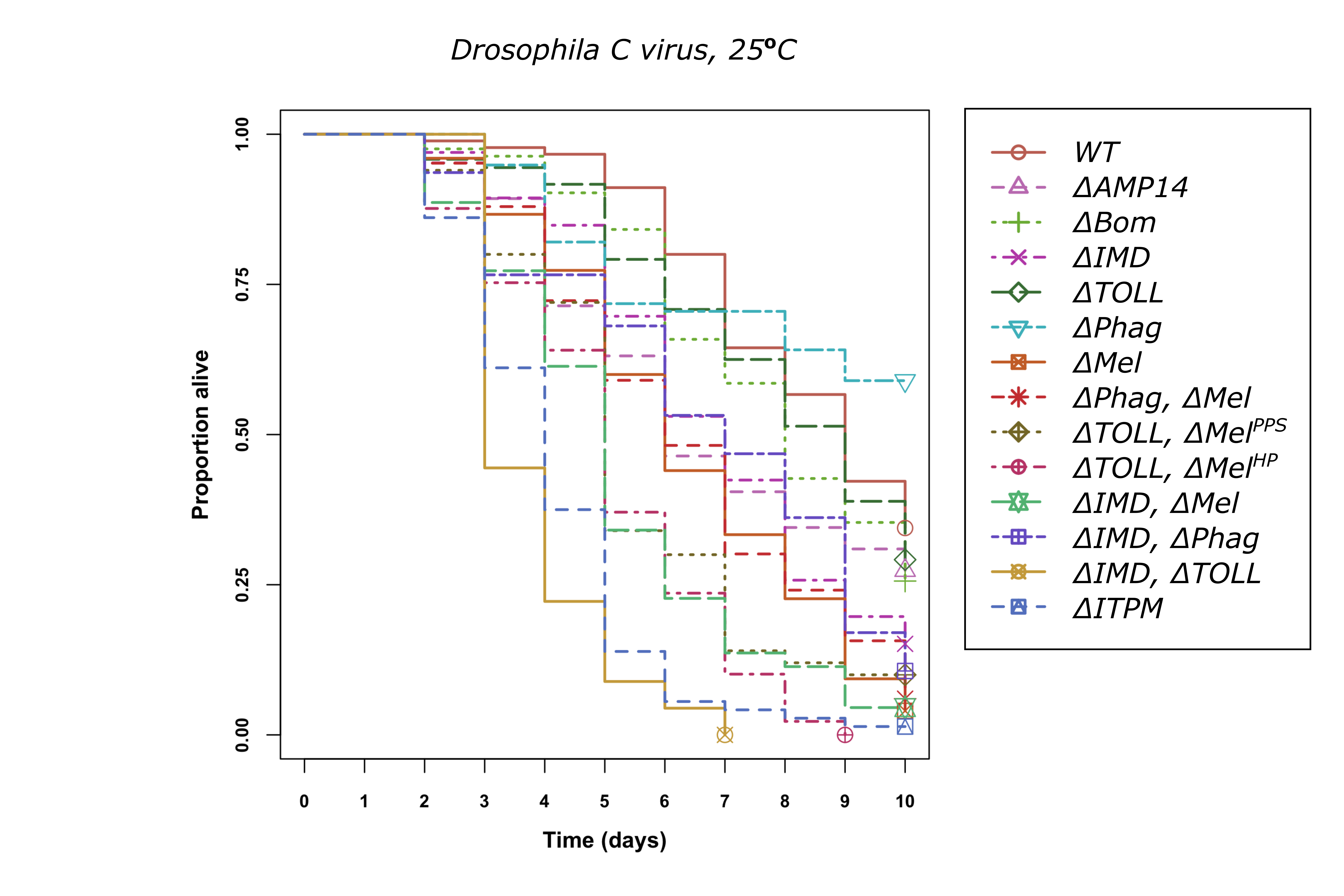

### DXV.png

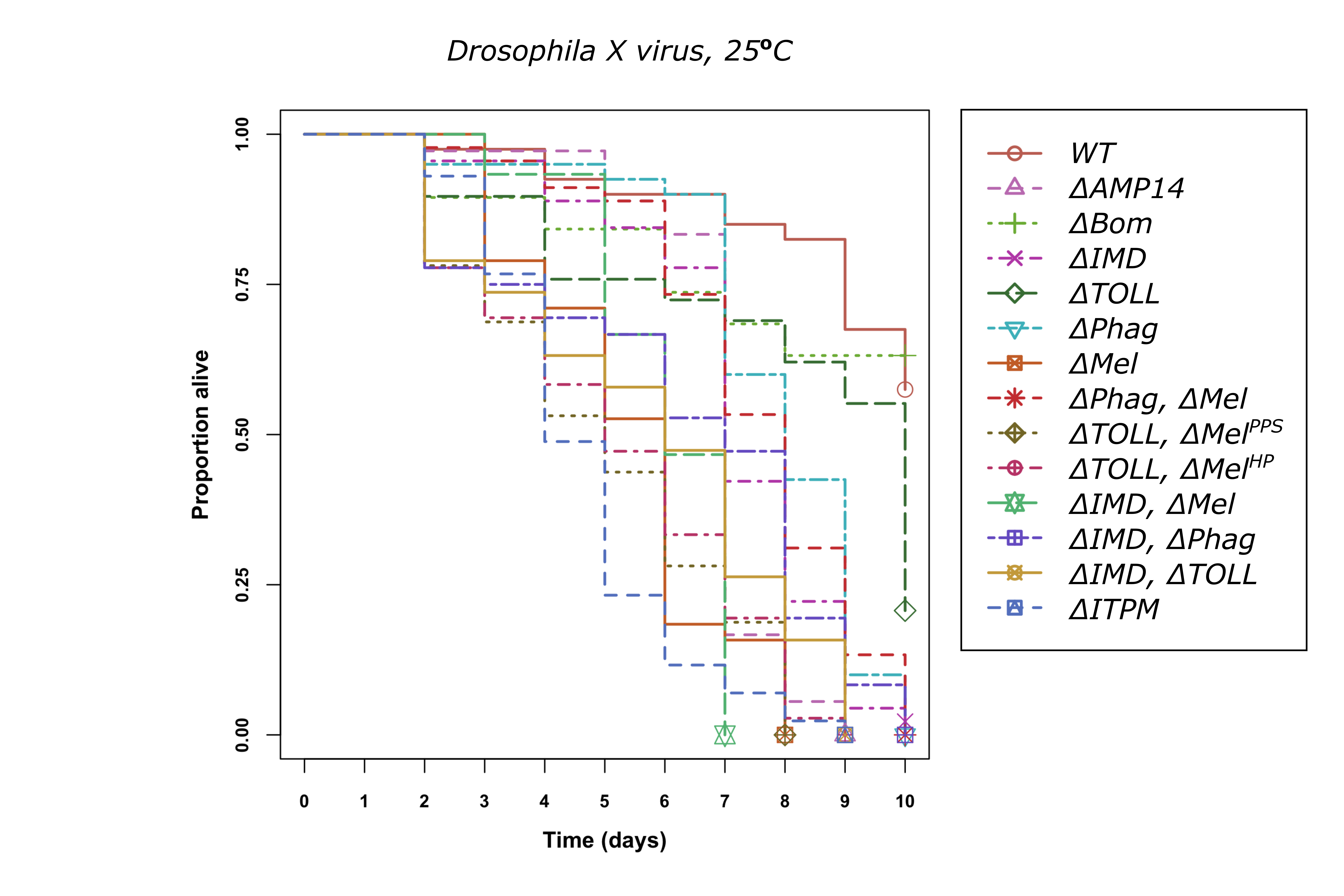

### E. cloacae.png

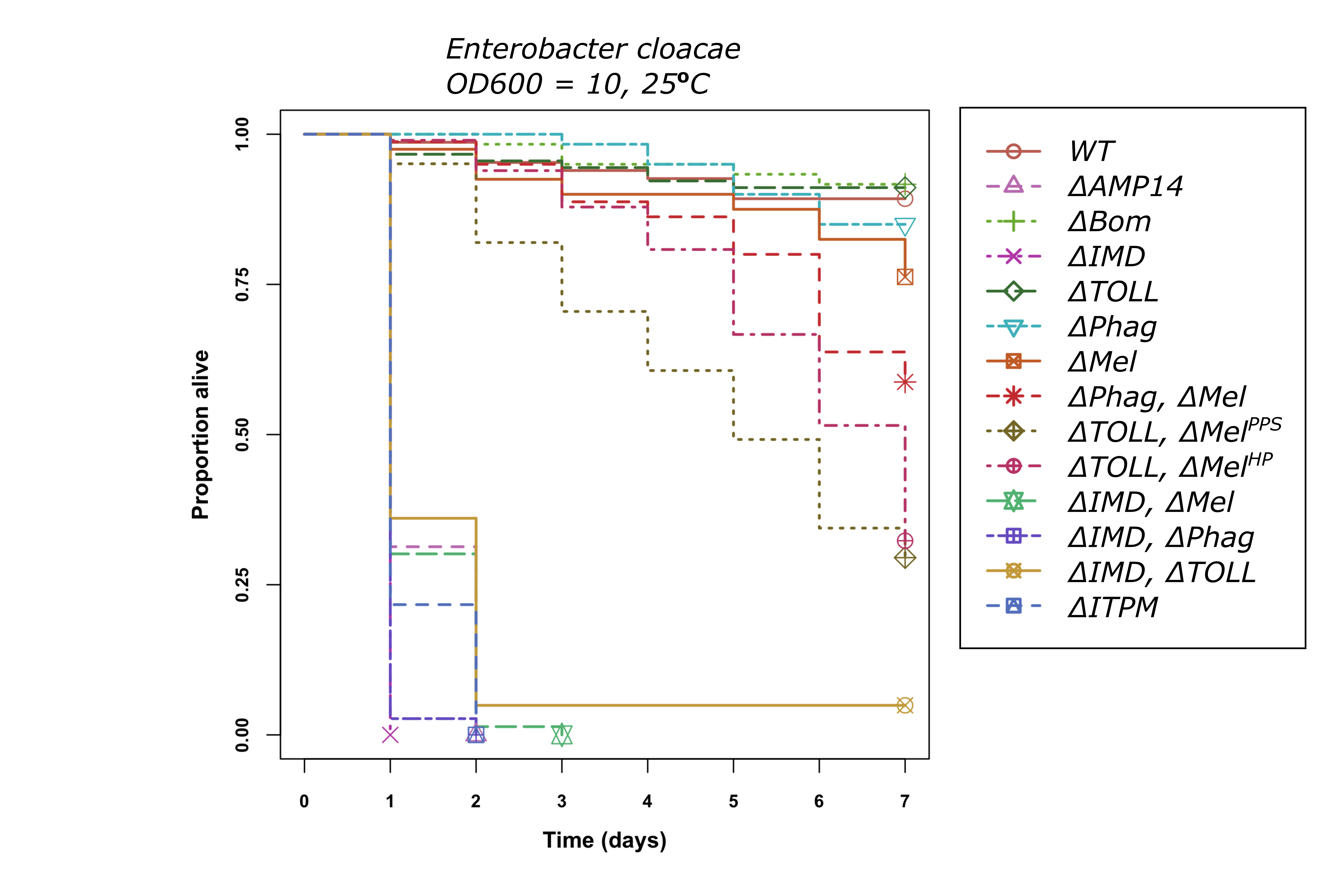

### E. coli.png

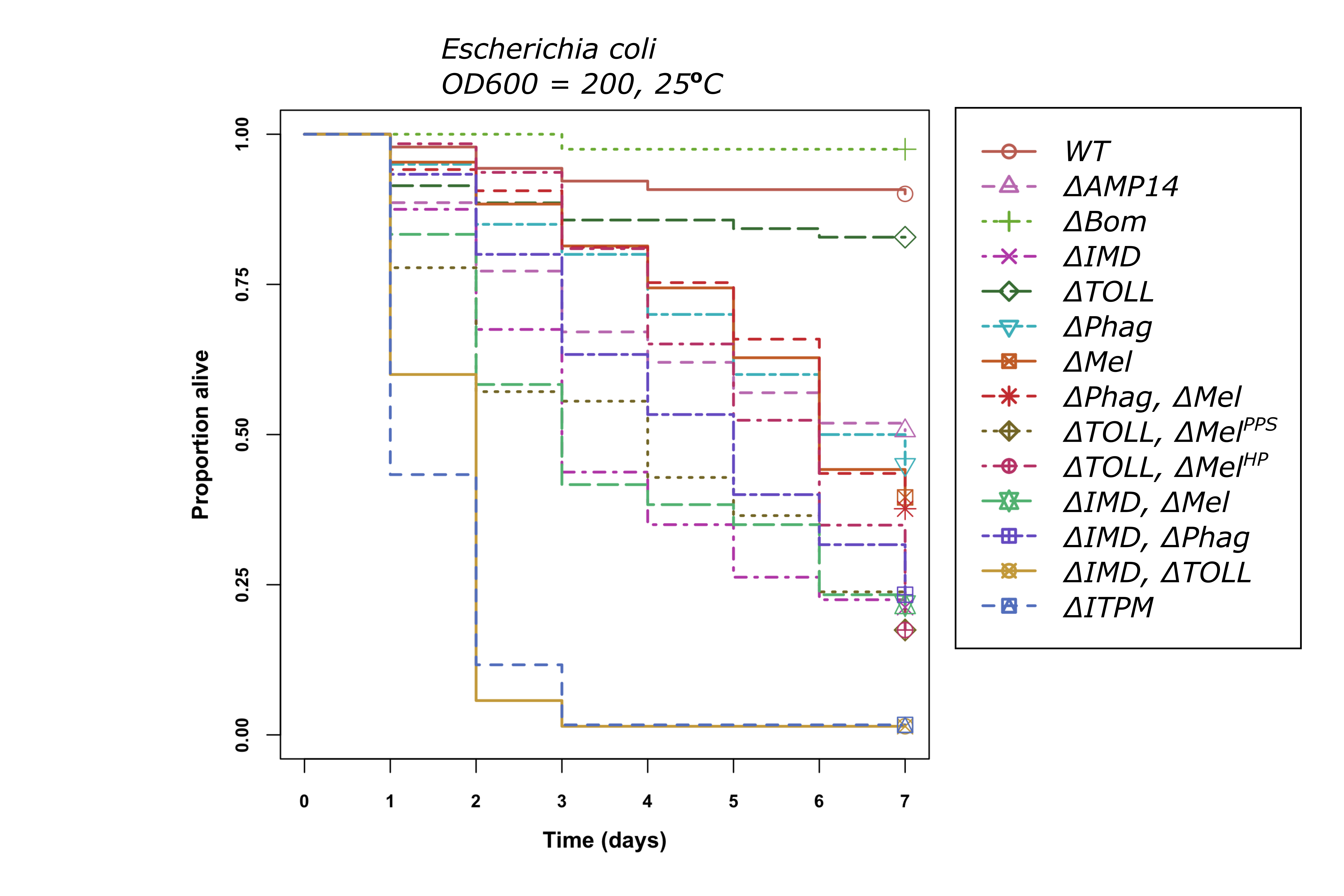

### E. faecalis.png

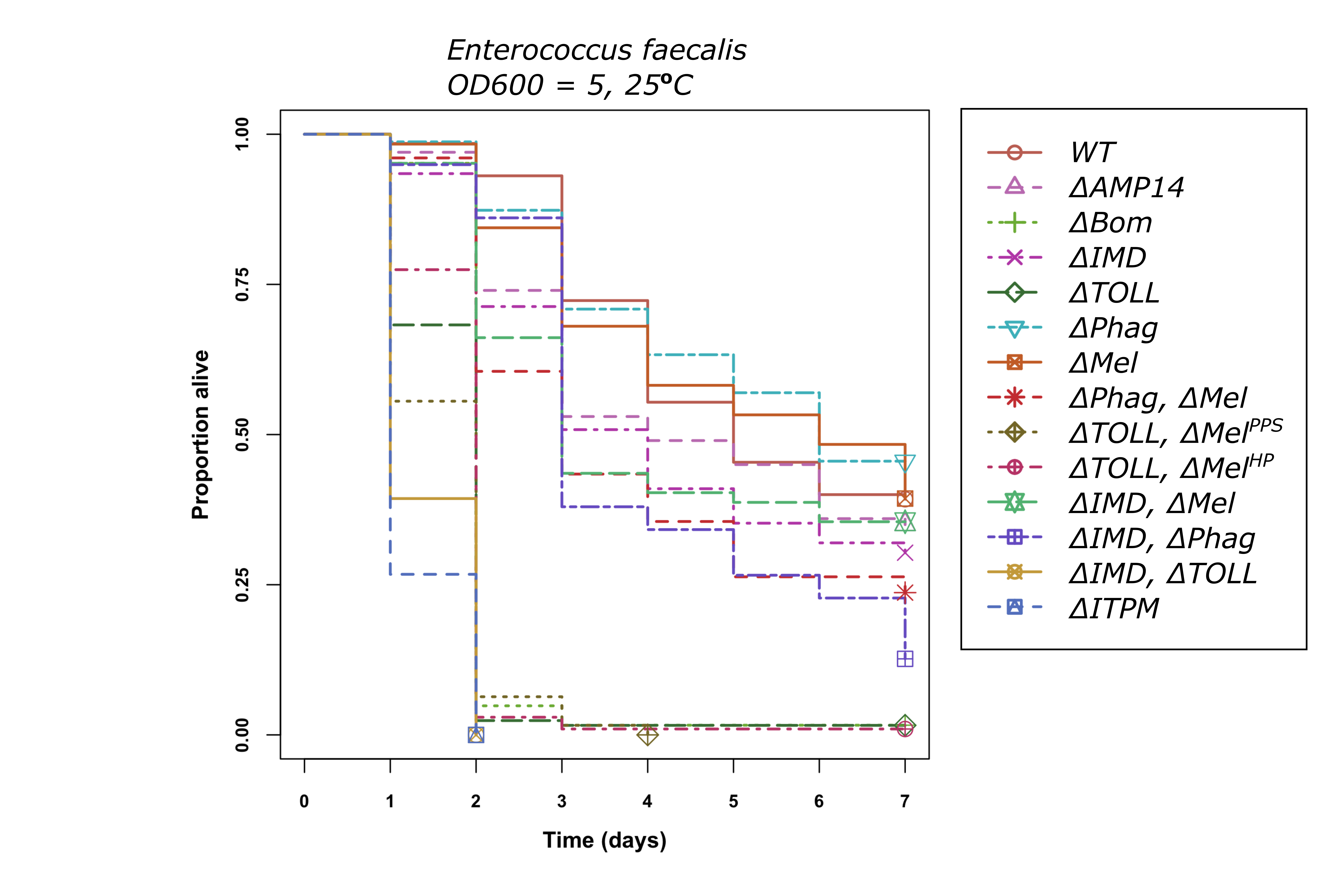

### FHV.png

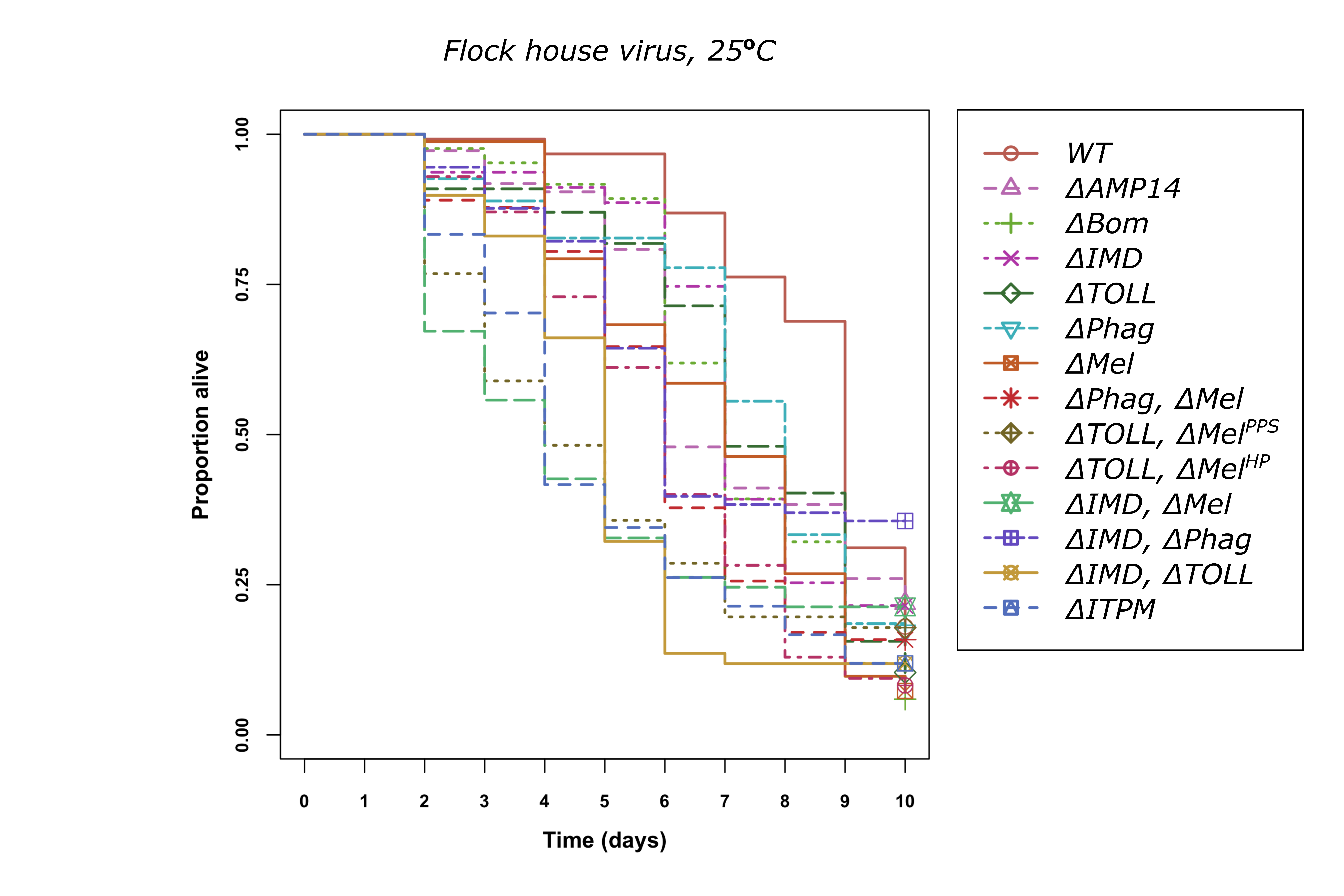

### IIV6.png

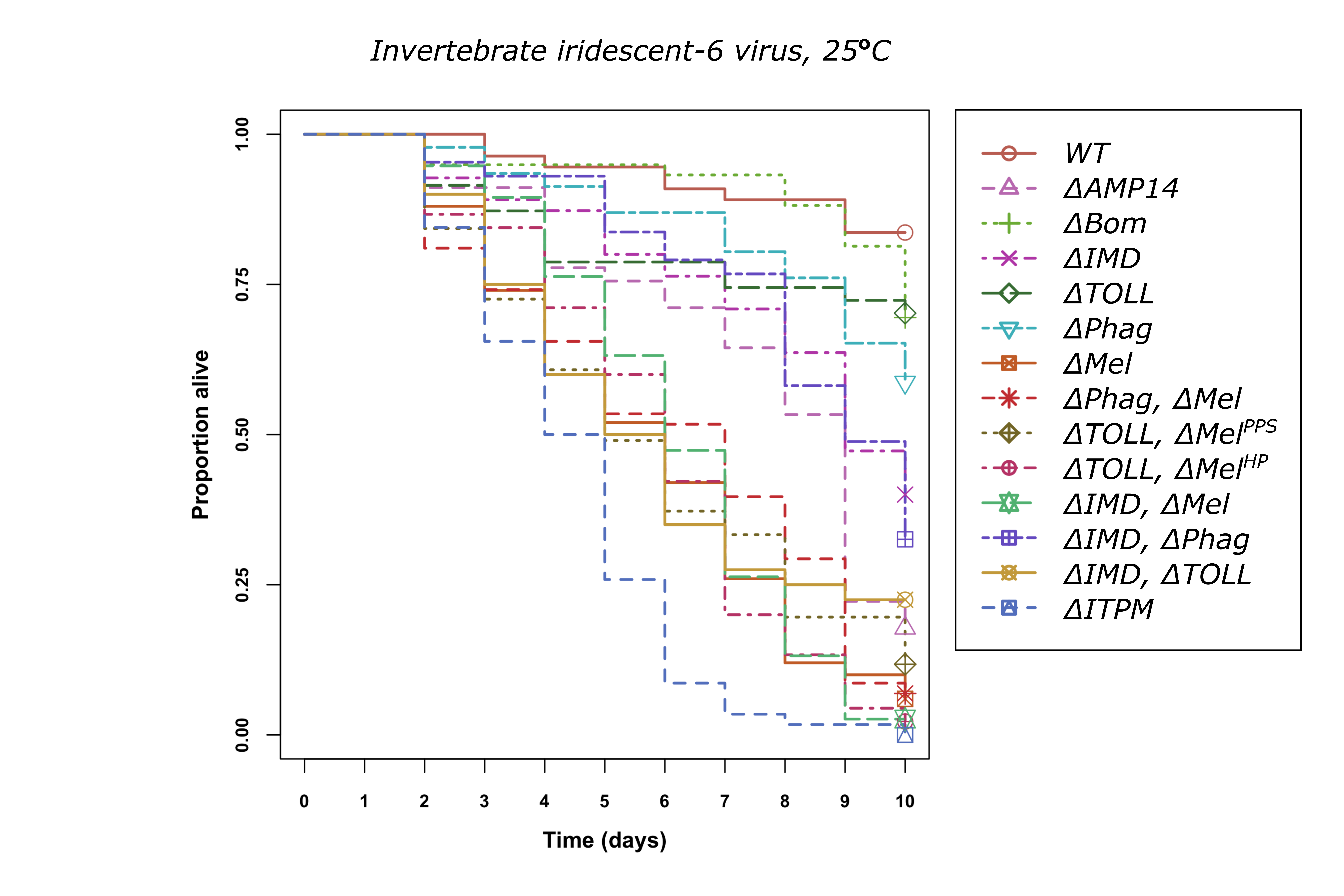

### K. pneumoniae.png

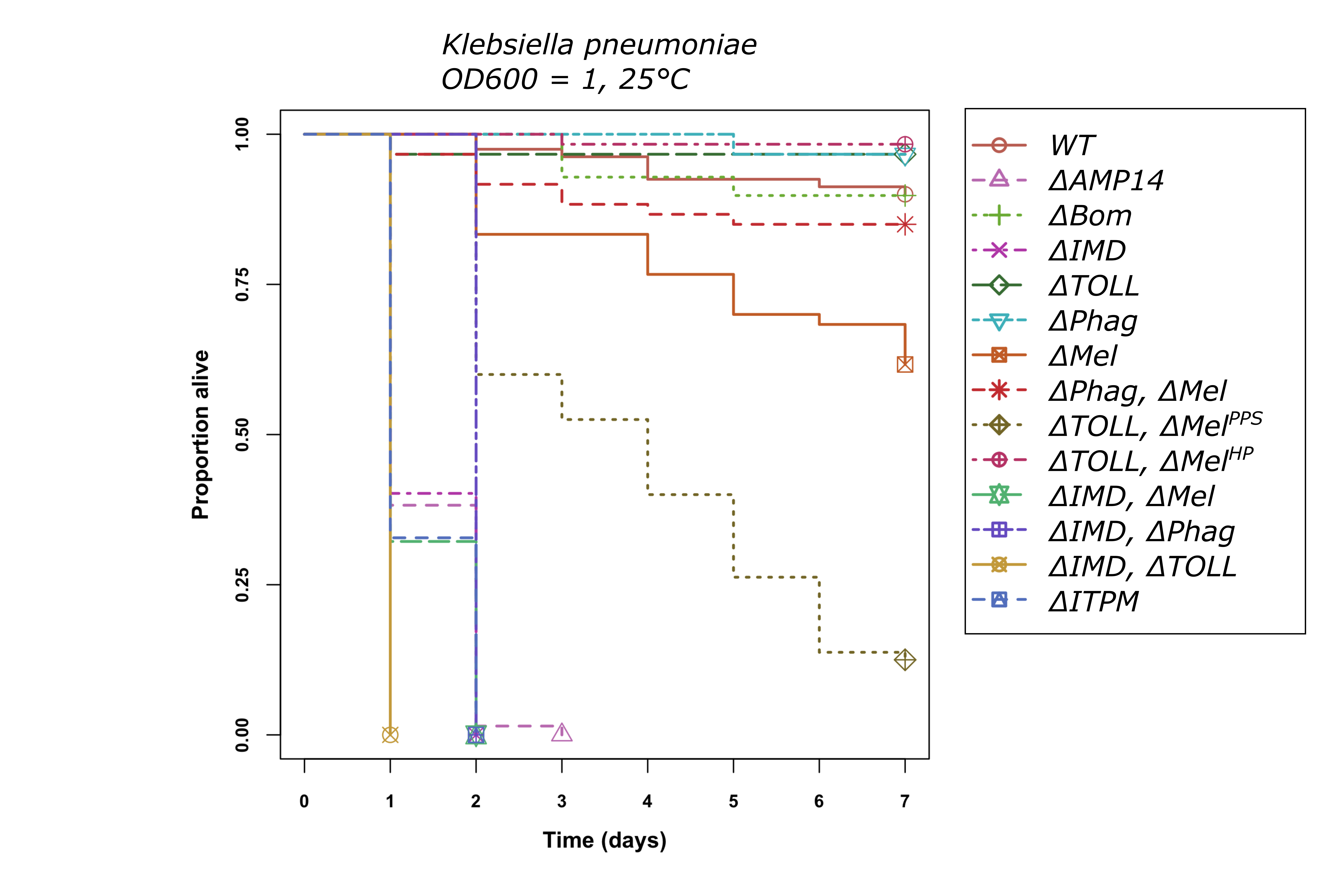

### L. monocytogenes.png

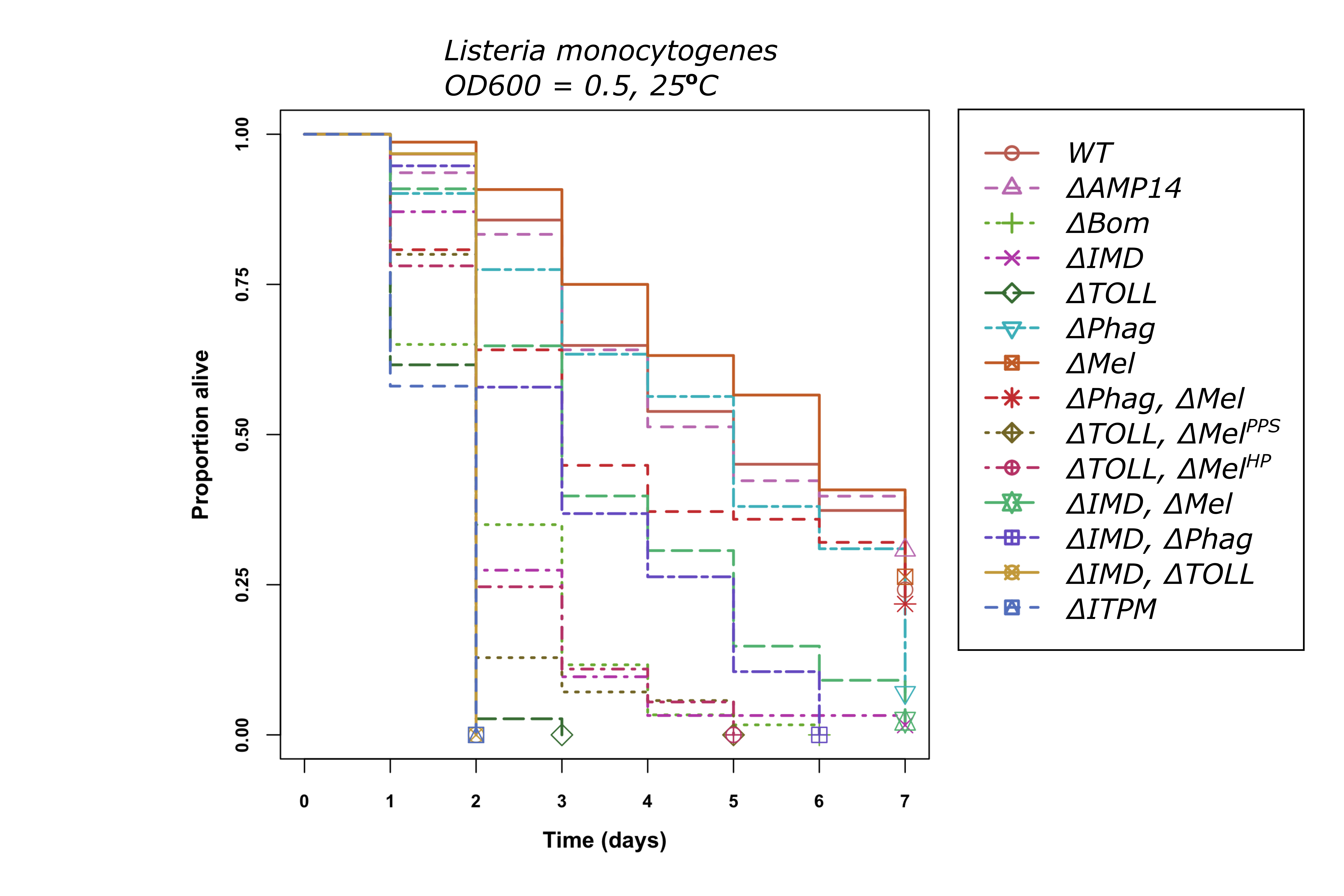

### Mi. luteus.png

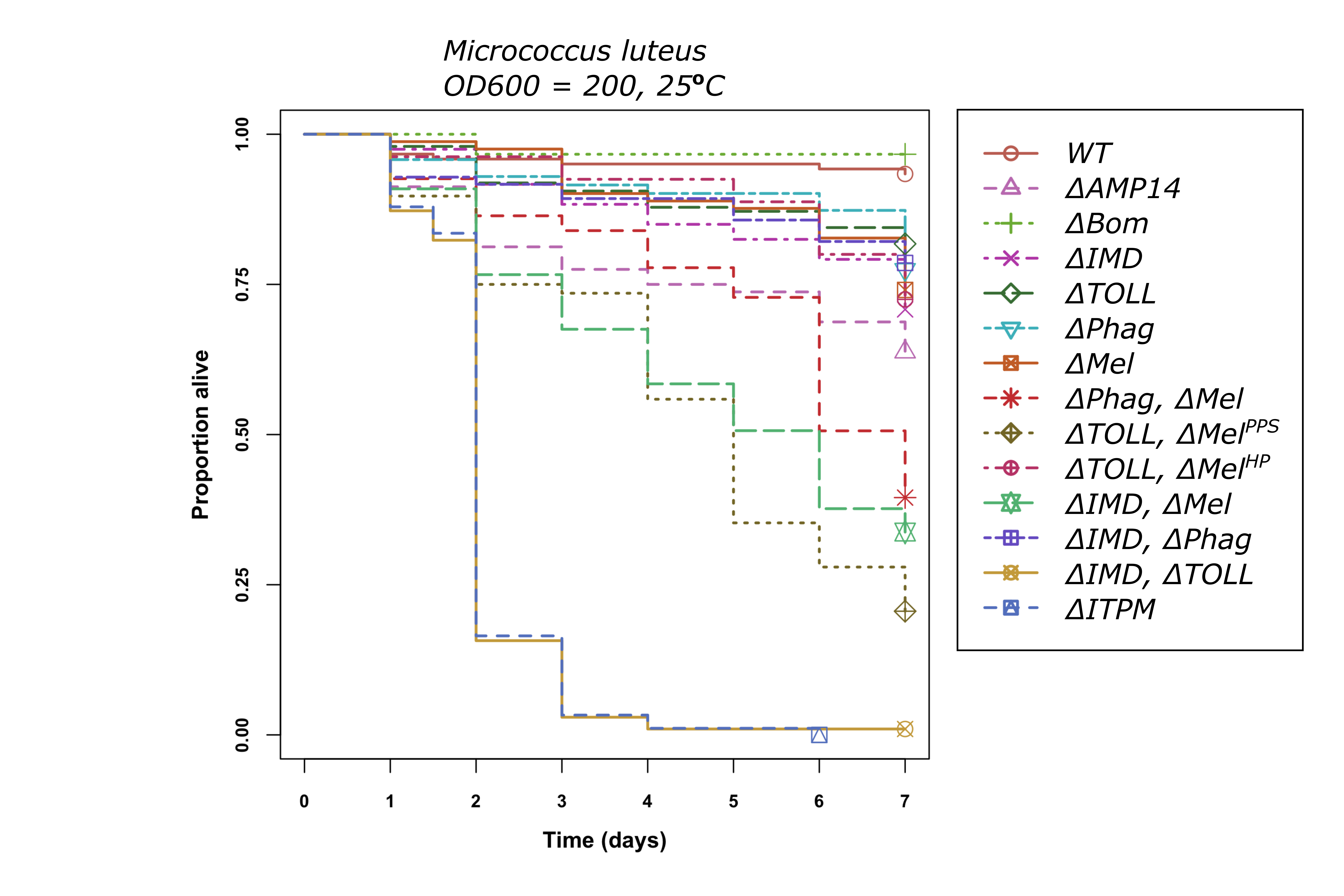

### My. marinum.png

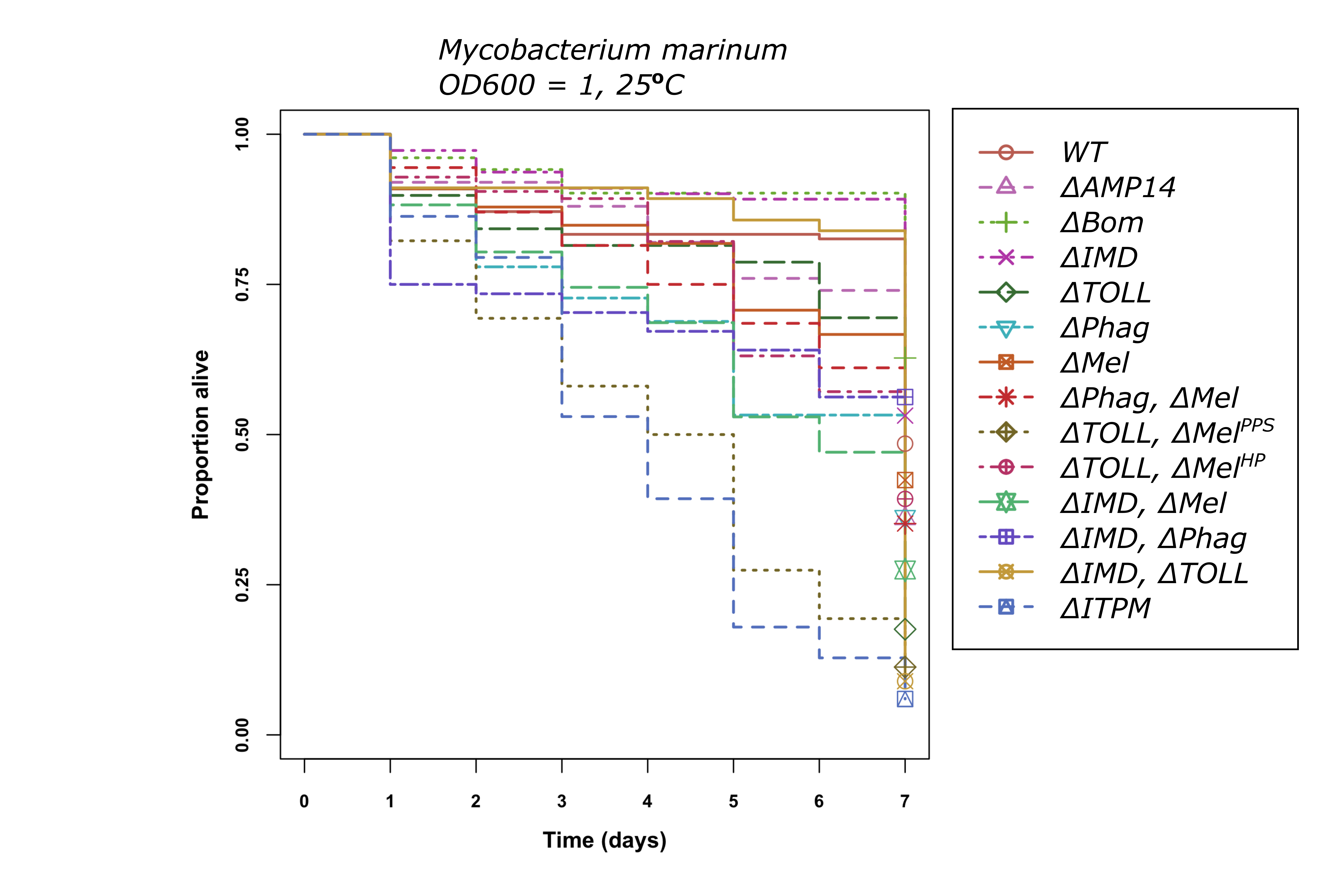

### Pe. carotovorum Ecc15.png

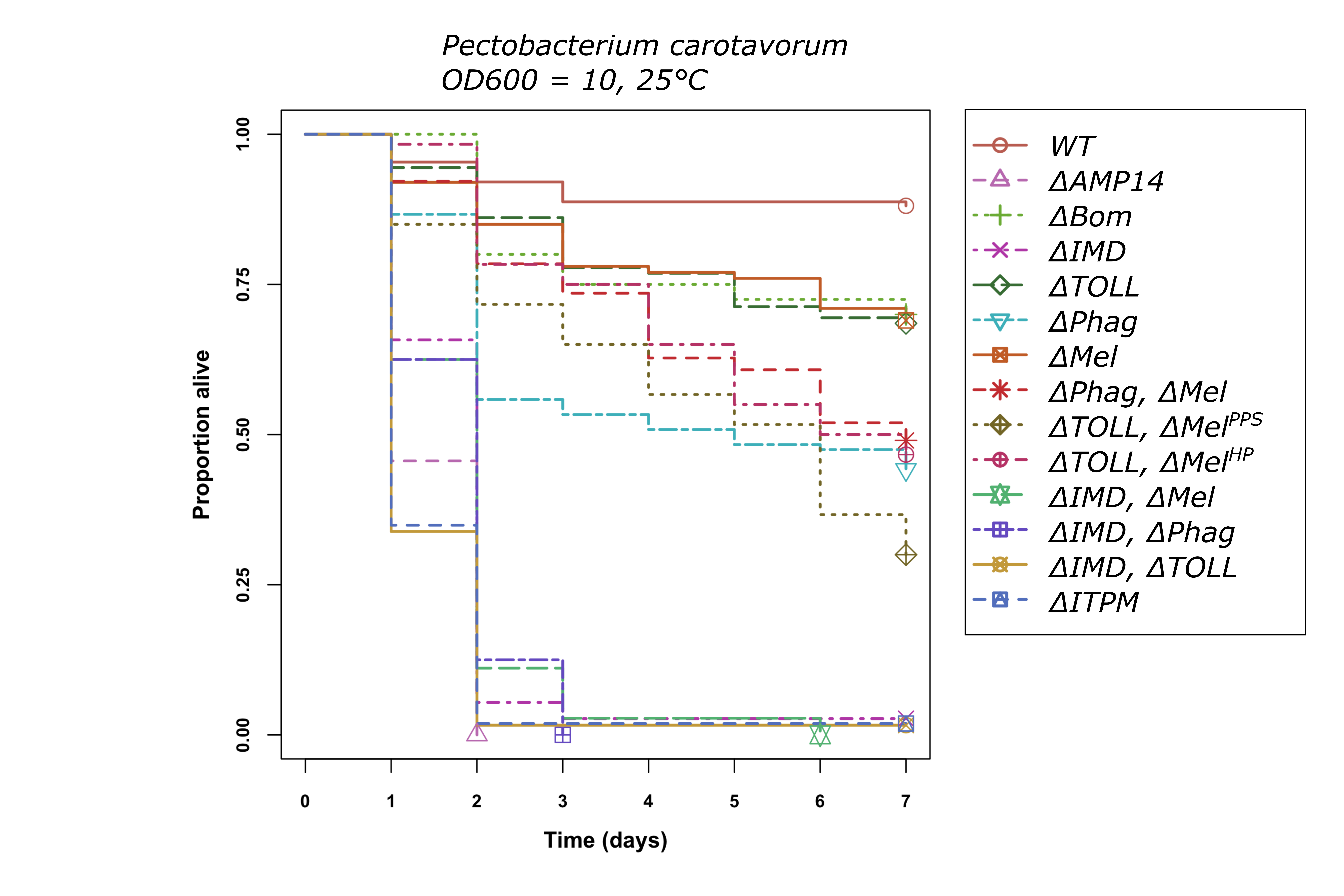

### Pr. burhodogranariea.png

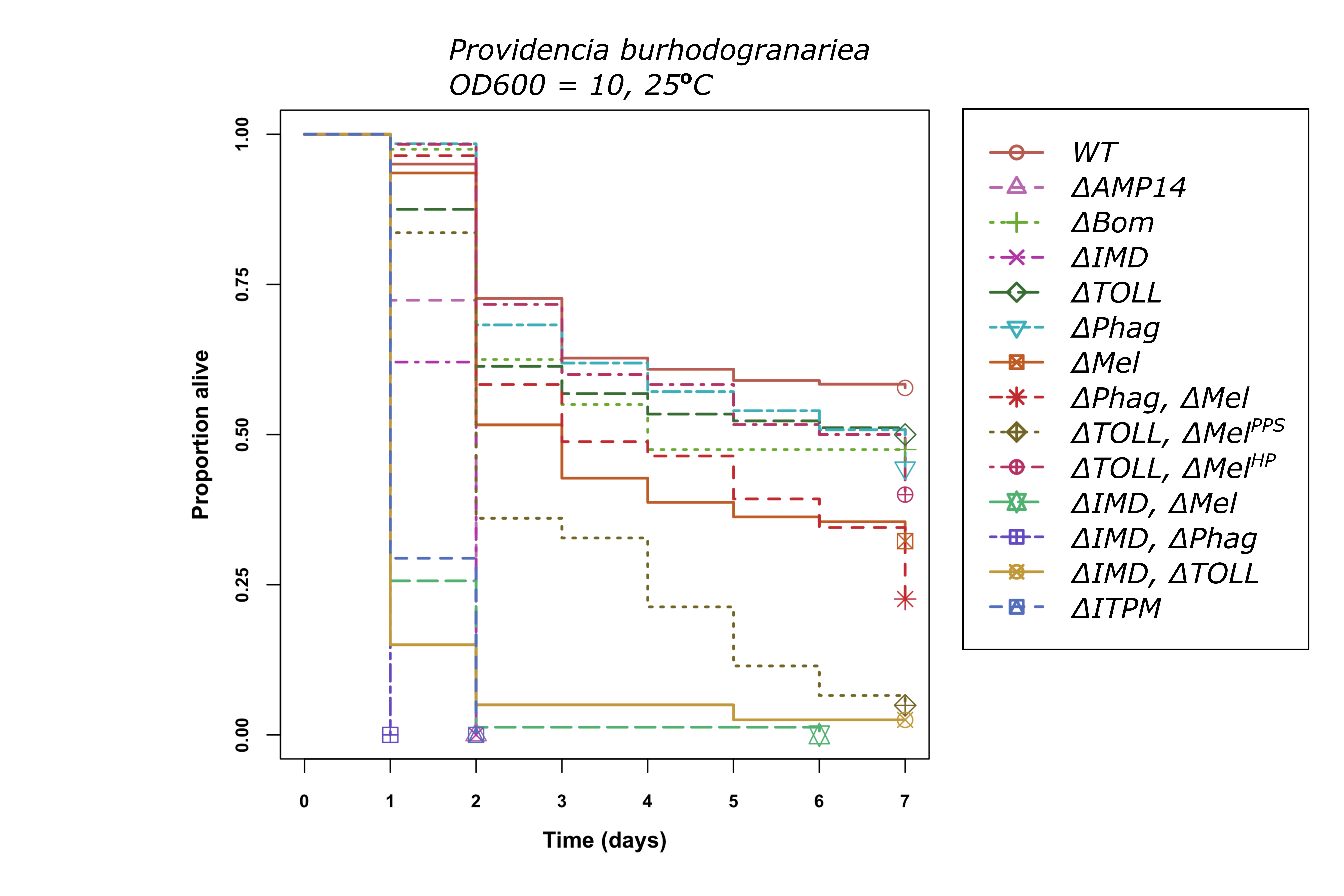

### Pr. rettgeri.png

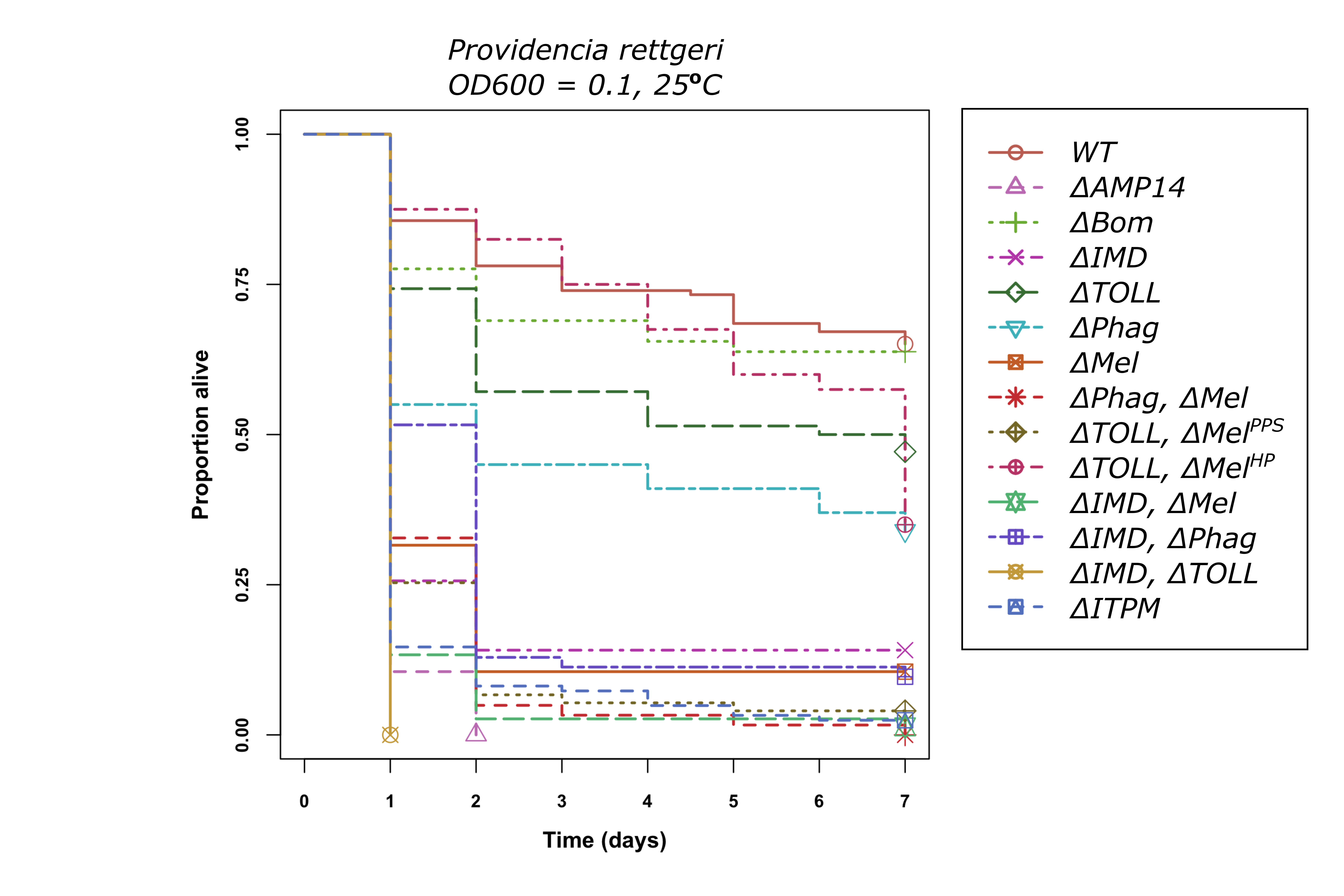

### Sa. typhimurium.png

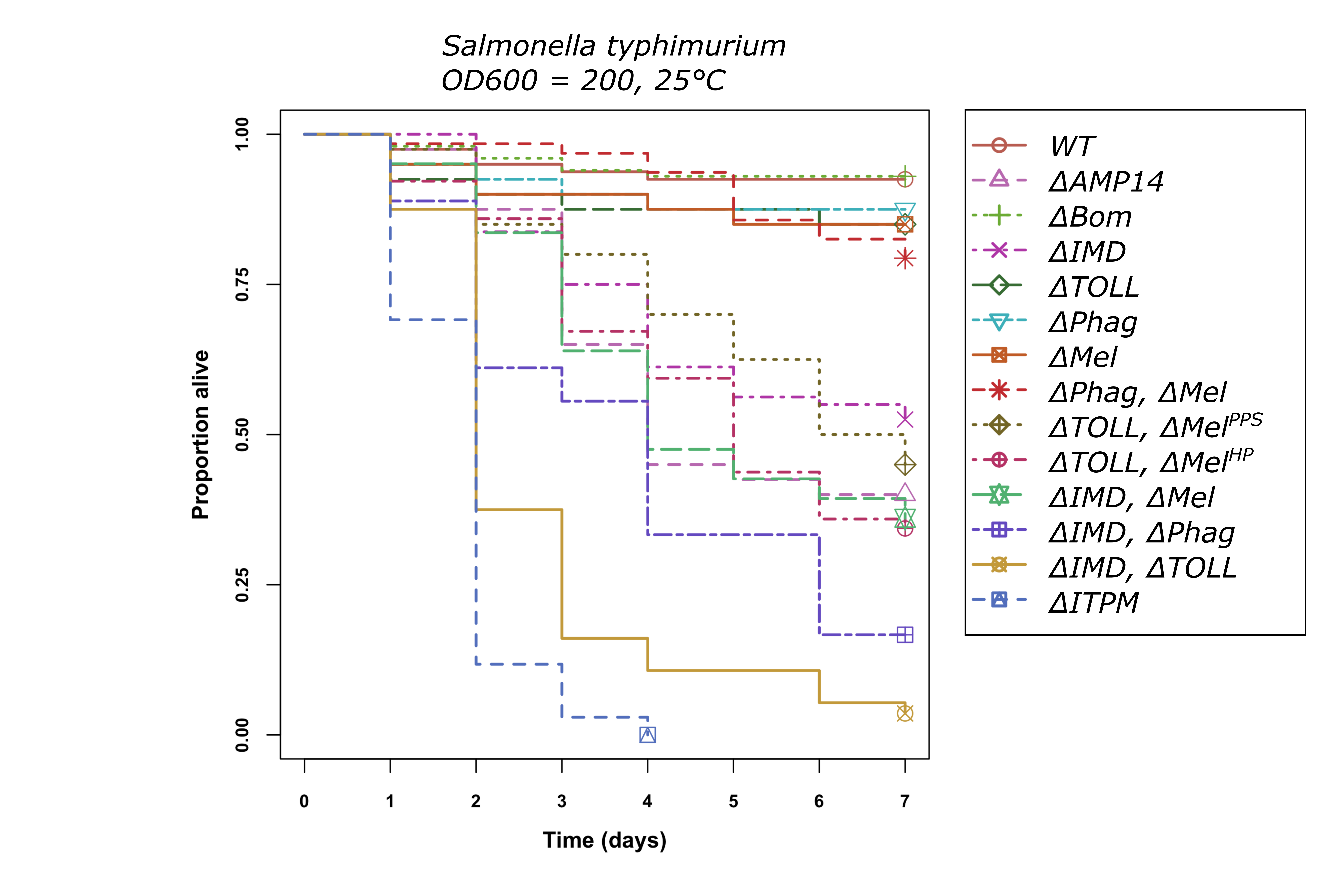

### SINV.png

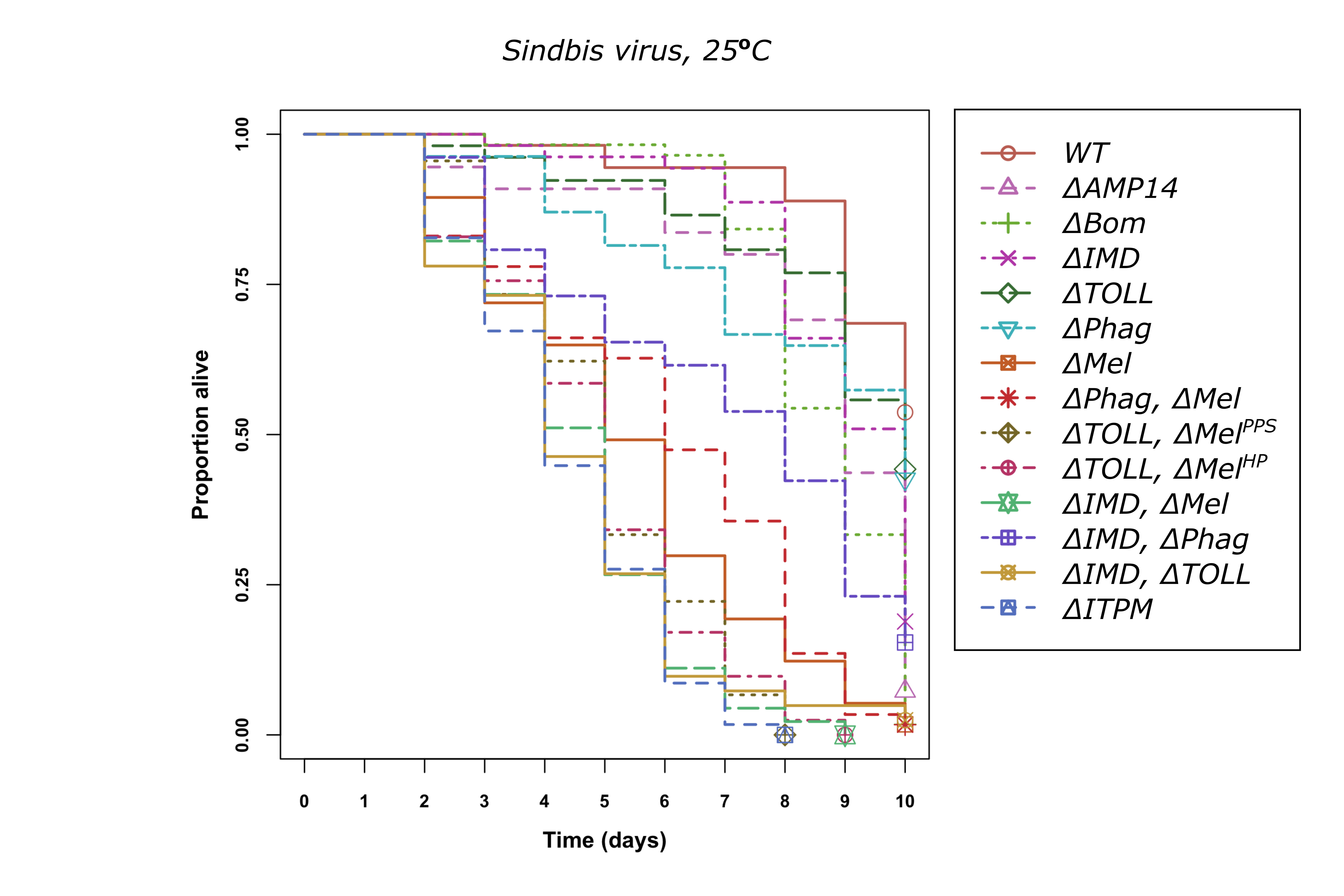

### Sta. aureus.png

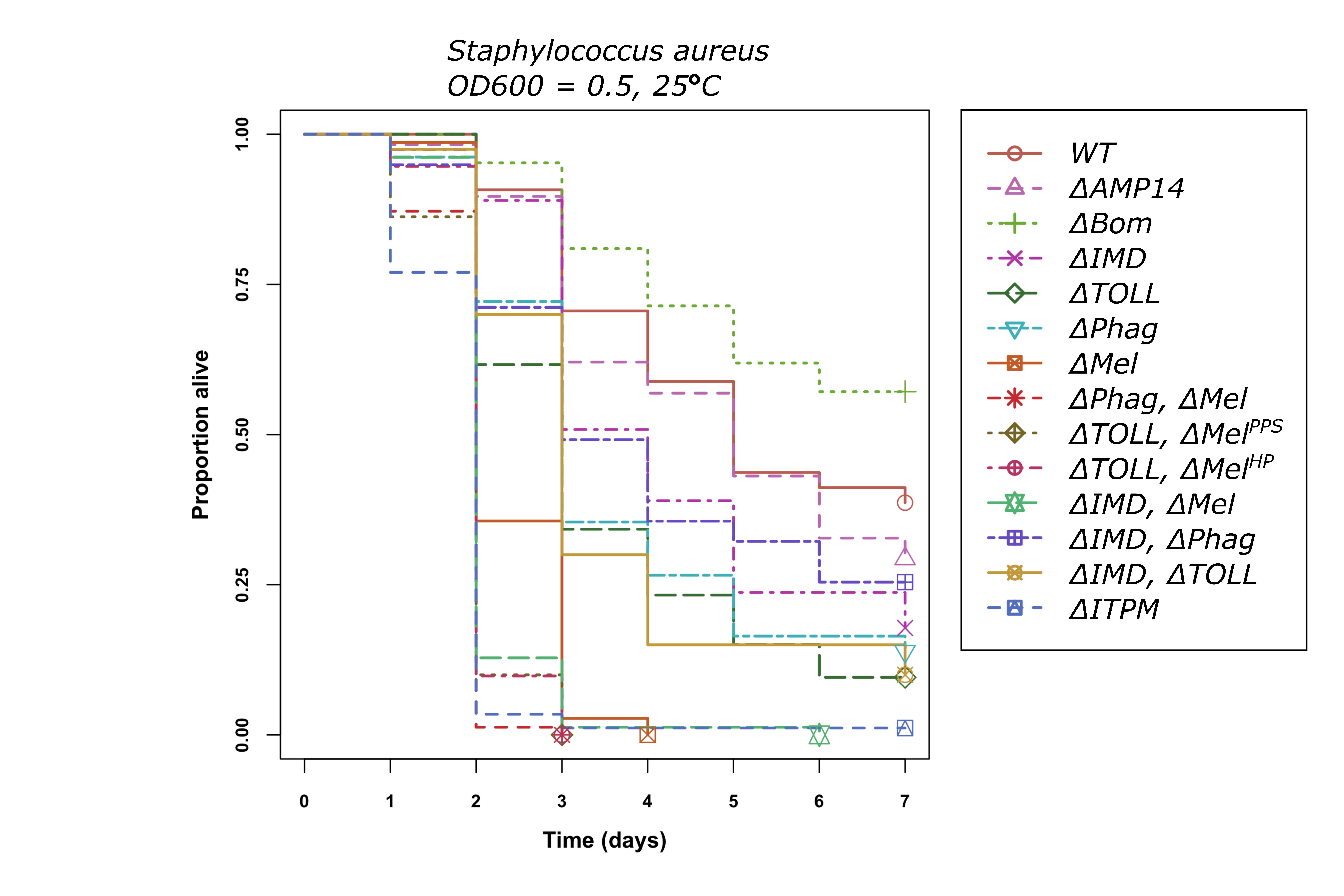

### Str. pneumoniae.png

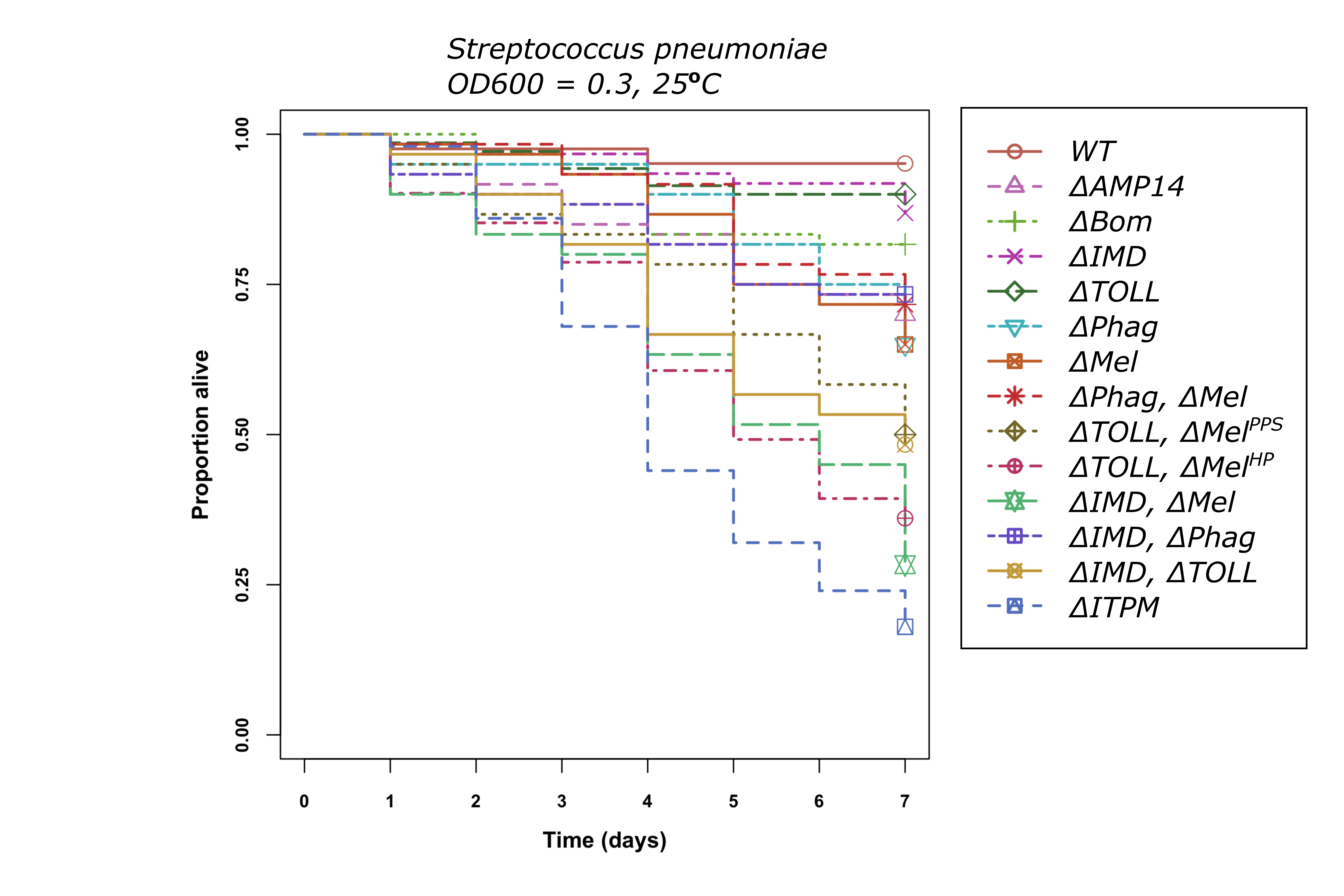

### V. parahemolyticus.png

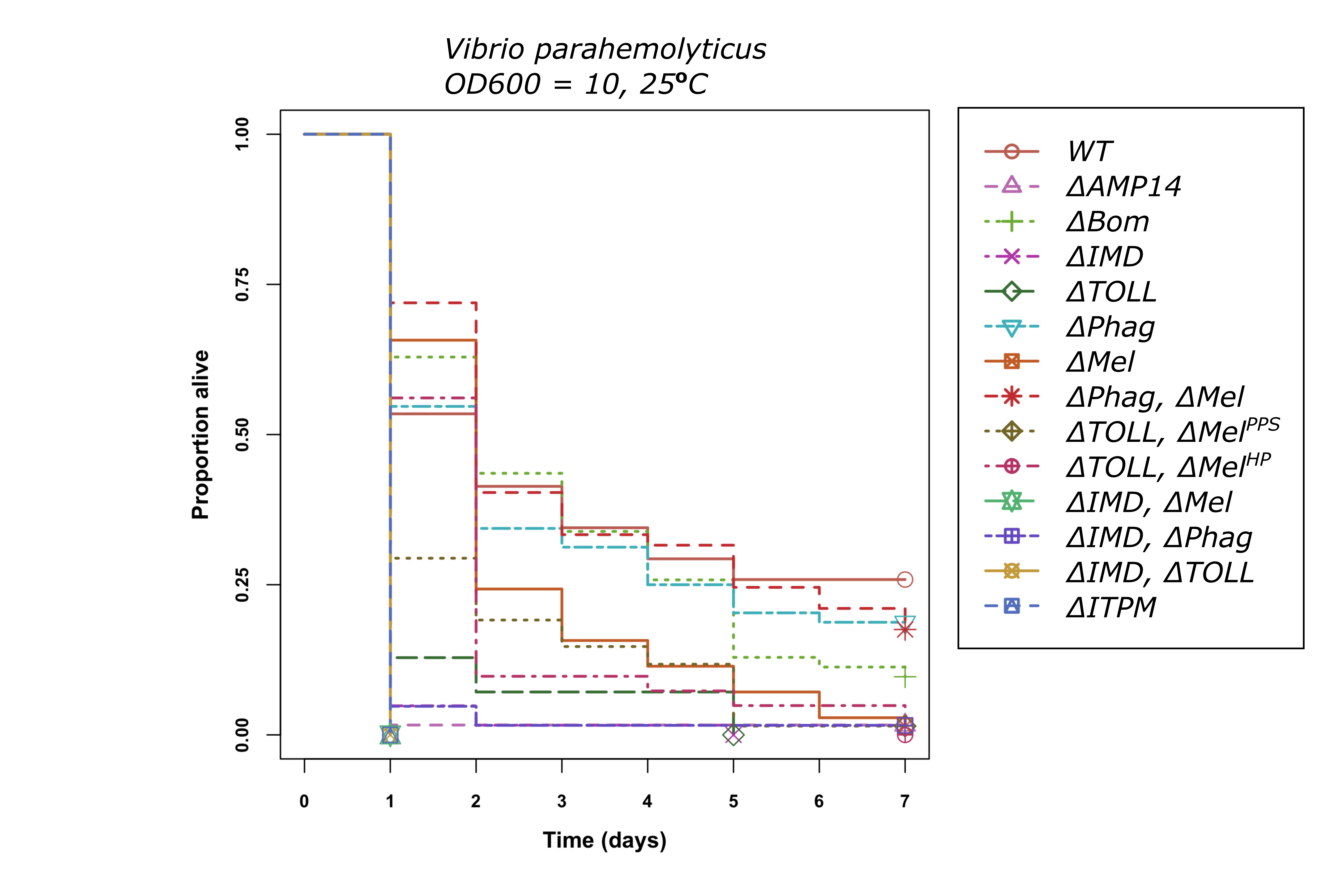
