## Supplementary material for "Layers of immunity: Deconstructing the *Drosophila* effector response": Figure 3supp1

| | **Microbe** |
| --- | --- |
| **Viruses** | DCV |
| | FHV |
| | DXV |
| | IIV-6 |
| | SINV |
| **Fungi** | *A. fumigatus (NI)* |
| | *B. bassiana (NI)* |
| | *B. bassiana (SI)* |
| | *C. albicans (SI)* |
| **Gram positive bacteria** | *M. marinum* |
| | *C. diphteriae* |
| | *M. luteus* |
| | *S. pneumoniae* |
| | *E. faecalis* |
| | *S. aureus* |
| | *L. monocytogenes* |
| | *B. subtilis* |
| **Gram negative bacteria** | *V. parahemolyticus* |
| | *P. rettgeri* |
| | *P. burhodogranaria* |
| | *Ecc15* |
| | *K. pneumoniae* |
| | *E. cloacae* |
| | *E. coli* |
| | *S. enterica typhimurium* |

| *AMP14* | *Bom* |
|---|---|
| ✓ | [redacted] |
| ✓ | ✓ |
| ✓ | [redacted] |
| ✓ | [redacted] |
| [redacted] | [redacted] |
| [redacted] | [redacted] |
| ✓ | [redacted] |
| [redacted] | ✓ |
| ✓ | [redacted] |
| [redacted] | [redacted] |
| [redacted] | [redacted] |
| [redacted] | [redacted] |
| [redacted] | [redacted] |
| [redacted] | ✓ |
| [redacted] | [redacted] |
| [redacted] | ✓ |
| ✓ | ✓ |
| ✓ | [redacted] |
| ✓ | [redacted] |
| ✓ | [redacted] |
| ✓ | ✓ |
| ✓ | [redacted] |
| ✓ | [redacted] |
| ✓ | [redacted] |
| ✓ | [redacted] |

| IMD | TOLL |
|---|---|
| ✓ | (✓ IM) |
| ✓ | ✓ |
| ✓ | ✓ |
| ✓ | (✓ I) |
| (✓ T, P) | (✓ I) |
| (✓ T) | (✓ I, M) |
| (✓ P) | (✓ I, M) |
| [redacted] | ✓ |
| (✓ T) | ✓ |
| [redacted] | [redacted] |
| (✓ T) | (✓ I, M) |
| (✓ T) | (✓ I) |
| [redacted] | [redacted] |
| | ✓ |
| | ✓ |
| ✓ | ✓ |
| ✓ | ✓ |
| ✓ | ✓ |
| ✓ | ✓ |
| ✓ | (✓ M) |
| ✓ | [redacted] |
| ✓ | |
| ✓ | |
| ✓ | (✓ I) |
| ✓ | (✓ I) |

| Phag | Mel |
|---|---|
| [redacted] | ✓ |
| ✓ | ✓ |
| ✓ | ✓ |
| [redacted] | ✓ |
| ✓ | ✓ |
| [redacted] | (✓ T) |
| (✓ I, M) | (✓ T, P) |
| ✓ | [redacted] |
| [redacted] | (✓ T) |

| | |
|---|---|
| [redacted] | [redacted] |
| [redacted] | ✓ |
| [redacted] | [redacted] |
| (✓ M) | (✓ P) |
| ✓ | ✓ |
| [redacted] | [redacted] |
| [redacted] | (✓ T) |

| | |
|---|---|
| [redacted] | [redacted] |
| ✓ | ✓ |
| [redacted] | ✓ |
| ✓ | [redacted] |
| [redacted] | [redacted] |
| ✓ | [redacted] |
| (✓ I) | [redacted] |
