## Supplementary material for "Layers of immunity: Deconstructing the *Drosophila* effector response": Table S1

| Type | Kingdom | Phylum | Class | Order | Family |
|---|---|---|---|---|---|
| Viruses | Bamfordvirae | Nucleocytoviricota | Megaviricetes | Pimascovirales | Iridoviridae |
| | Orthornavirae | nd | nd | nd | Birnaviridae |
| | Orthornavirae | Pisuviricota | Pisoniviricetes | Picornavirales | Dicistroviridae |
| | Orthornavirae | Kitrinoviricota | Magsaviricetes | Nodamuvirales | Nodaviridae |
| | Orthornavirae | Kitrinoviricota | Alsuviricetes | Martelliirales | Togaviridae |
| Fungi | Fungi | Ascomycota | Sacharomycete | Sacharomycetales | Sacharomycetacae |
| | | Ascomycota | Eurotiomycete | Eurotiales | Trychocomacae |
| | | Ascomycota | Sordaryomycete | Coricypitacae | Ophiocordycipitacae |
| Gram positive bacteria | Bacteria | Actinomycetota | Actinomycetia | Mycobacteriales | Mycobacteriacae |
| | | Actinomycetota | Actinomycetia | Mycobacteriales | Corynebacteriaceae |
| | | Actinomycetota | Actinomycetia | Actinomycetales | Micrococcacae |
| | | Bacillota | Bacilli | Lactobacilliales | Streptococcacae |
| | | Bacillota | Bacilli | Lactobacilliales | Enterococcacae |
| | | Bacillota | Bacilli | Caryophanale | Staphylococcacae |
| | | Bacillota | Bacilli | Bacilliales | Listeriacae |
| | | Bacillota | Bacilli | Bacilliales | Bacillacae |
| Gram negative bacteria | Bacteria | Pseudomonadota | Gammaproteobacteriacae | Pseudomonadales | Pseudomonadacae |
| | | Pseudomonadota | Gammaproteobacteriacae | Vibrionales | Vibrionacae |
| | | Pseudomonadota | Gammaproteobacteriacae | Enterobacteriales | Morgenallacae |
| | | Pseudomonadota | Gammaproteobacteriacae | Enterobacteriales | Morgenallacae |
| | | Pseudomonadota | Gammaproteobacteriacae | Enterobacteriales | Enterobacteriacae |
| | | Pseudomonadota | Gammaproteobacteriacae | Enterobacteriales | Enterobacteriacae |
| | | Pseudomonadota | Gammaproteobacteriacae | Enterobacteriales | Enterobacteriacae |
| | | Pseudomonadota | Gammaproteobacteriacae | Enterobacteriales | Enterobacteriacae |
| | | Pseudomonadota | Gammaproteobacteriacae | Enterobacteriales | Enterobacteriacae |

| Nucleic acid / Membrane | Species | Strain, Source, DOI of previous study | Inf. mode | Infection temperature | Used concentration | Growth temperature |
|---|---|---|---|---|---|---|
| dsDNA+ | ***Invertebrate iridescent-6 virus*** | gift from Ronald van Rij | intra-thoracique | 25°C | 31 700 TCID50 | provided |
| ssRNA+ | ***Drosophila X virus*** | gift from Ronald van Rij | intra-thoracique | 25°C | 431 000 TCID50 | provided |
| ssRNA+ | ***Drosophila C virus*** | gift from Carla Saleh | intra-thoracique | 25°C | 2 000 TCID50 | 26°C |
| dsRNA+ | ***Flock House virus*** | gift from Carla Saleh | intra-thoracique | 25°C | 250 000 TCID50 | 26°C |
| ssRNA+ | ***Sindbis virus*** | gift from Ronald van Rij | intra-thoracique | 25°C | 50 000 TCID50 | provided |
| *Beta-glucans* | **Candida albicans** | ATCC 2001 | septic | 29°C | $OD^{600}$= 200 | 37°C |
| *Beta-glucans* | ***Aspergillus fumigatus*** | *doi: 10.7554/eLife.44341* | natural | 29°C | rolled on plate | 37°C |
| *Beta-glucans* | ***Beauvaria bassiana*** | R444 | septic | 29°C | $OD^{600}$= 5 | commercial |
| *Beta-glucans* | | | natural | 29°C | 30mg | commercial |
| mAGP-type | ***Mycobacterium marinum*** | 1218R WT | intra-thoracique | 25°C | $OD^{600}$= 1 | 37°C |
| Lys-type | ***Corynebacterium diphtheriae*** | *DSM 44123* | septic | 25°C | $OD^{600}$= 100 | 37°C |
| Lys-type | ***Micrococcus luteus*** | *doi: 10.7554/eLife.44341* | septic | 25°C | $OD^{600}$= 200 | 29°C |
| Lys-type | ***Streptococcus pneumoniae*** | D39V WT | septic | 25°C | $OD^{600}$= 0.3 | 37°C |
| Lys-type | ***Entecoccus faecalis*** | 254 | septic | 25°C | $OD^{600}$= 5 | 37°C |
| Lys-type | ***Staphylococcus aureus*** | *doi: 10.7554/eLife.44341* | septic | 25°C | $OD^{600}$= 0.5 | 37°C |
| DAP-type | ***Listeria monocytogenes*** | BUG 1600 | septic | 25°C | $OD^{600}$= 0.5 | 37°C |
| DAP-type | ***Bacillus subtilis*** | 168 | septic | 25°C | $OD^{600}$= 5 | 37°C |
| DAP-type | ***Pseudomonas aeruginosa*** | PAO1 | septic | 25°C | $OD^{600}$= 0.01 | 37°C |
| DAP-type | ***Vibrio parahemolyticus*** | | septic | 25°C | $OD^{600}$= 10 | 37°C |
| DAP-type | ***Providencia rettgeri*** | Dmel | septic | 25°C | $OD^{600}$= 0.1 | 37°C |
| DAP-type | ***Providencia burhodogranaria*** | B | septic | 25°C | $OD^{600}$= 10 | 37°C |
| DAP-type | ***Pectobacterium carotovorum*** | Ecc15 | septic | 25°C | $OD^{600}$= 10 | 37°C |
| DAP-type | ***Klebsiella pneumoniae*** | | septic | 25°C | $OD^{600}$= 1.0 | 37°C |
| DAP-type | ***Enterobacter cloacae*** | B12 | septic | 25°C | $OD^{600}$= 10 | 37°C |
| DAP-type | ***Escherichia coli*** | 1106 | septic | 25°C | $OD^{600}$= 200 | 37°C |
| DAP-type | ***Salmonella enterica ser. typhimurium*** | *doi: 10.7554/eLife.44341* | septic | 25°C | $OD^{600}$= 200 | 37°C |

| Medium of culture |
|---|
| provided |
| provided |
| S2 cells in Schneider |
| S2 cells in Schneider |
| provided |
| YPG |
| Malt-agar |
| commercial |
| commercial |
| LB |
| LB |
| LB |
| LB |
| LB |
| BHI |
| LB |
| BHI |
| LB |
| LB |
| LB |
| LB |
| LB |
| LB |
| LB |
| LB |
| LB |
